# Investigating Novel *Streptomyces* Bacteriophage Endolysins as Potential Antimicrobial Agents

**DOI:** 10.1101/2024.04.29.591658

**Authors:** Jindanuch Maneekul, Amanda Chiaha, Rachel Hughes, Faith Labry, Joshua Saito, Matthew Almendares, Brenda N. Banda, Leslie Lopez, Nyeomi McGaskey, Melizza Miranda, Jenil Rana, Brandon R. Zadeh, Lee E. Hughes

## Abstract

**Background:** As antibiotic resistance has become a major global threat; the World Health Organization (WHO) has urgently called for alternative strategies for control of bacterial infections. Endolysin, a phage-encoded protein, can degrade bacterial peptidoglycan (PG) and disrupt bacterial growth. According to the WHO, there are only three endolysin products currently in clinical phase development. In this study we explore novel endolysins from *Streptomyces* phages as only a few of them have been experimentally characterized. Using several bioinformatics tools, we identified nine different functional domain combinations from 250 *Streptomyces* phages putative endolysins. LazerLemon gp35 (CHAP; LL35lys), Nabi gp26 (amidase; Nb26lys), and Tribute gp42 (PGRP/amidase; Tb42lys) were selected for experimental studies. We hypothesized that (1) the proteins of interest will have the ability to degrade purified PG, and (2) the proteins will have potential antimicrobial activity against bacteria from families of importance in antibiotic resistance, such as ESKAPE safe relatives (*Enterococcus raffinosus, Staphylococcus epidermidis*, *Klebsiella aerogenes*, *Acinetobacter baylyi*, *Pseudomonas putida*, and *Escherichia coli*).

**Results:** LL35lys, Nb26lys, and Tb42lys exhibit PG-degrading activity on zymography and hydrolysis assay. The enzymes (100 µg/mL) can reduce PG turbidity to 32-40%. The killing assay suggests that Tb42lys has a broader range (*E. coli*, *P. putida*, *A. baylyi* and *K. aerogenes*). While Nb26lys better attacks Gram-negative than Gram-positive bacteria, LL35lys can only reduce the growth of the Gram-positive ESKAPE strains but does so effectively with a low MIC_90_ of 2 µg/mL. A higher concentration (≥300 µg/mL) of Nb26lys is needed to inhibit *P. putida* and *K. aerogenes*.

**Conclusion:** From 250 putative endolysins, bioinformatic methods were used to select three putative endolysins for cloning and study: LL35lys, Nb26lys, and Tb42lys. All have shown PG-degrading activity, a critical function of endolysin. With a low MIC, LL35lys shows activity for the Gram-positive ESKAPE strains, while Nb26lys and Tb42lys are active against the Gram-negatives. Therefore, endolysins from *Streptomyces* phage have potential as possible antimicrobial agents against ESKAPE bacteria.

## 1 Background

Lysin, a protein that can lyse the bacterial cell wall, has been intensively studied over the past couple of decades [1, 2] and has become a promising treatment for bacterial infections in recent years [3–5]. Endolysin is a lysin encoded by a gene from a bacteriophage (or phage for short). At a coordinated time in the phage infection cycle, endolysin breaks down the cell wall from inside, leading to cell death [1]. Peptidoglycan (PG) is a major component that forms a thick cell wall, located outside the cell membrane [6, 7]. The general molecular structure reveals that PG is a glycan tetrapeptide consisting of N-acetylglucosamine (NAG), N-acetylmuramic acid (NAM), and oligopeptide (L-ala, D-glu, *m*DAP or L-lys, D-ala, and D-ala, respectively) connected with the NAM to form a net-like structure. Degradation of peptidoglycan has been extensively studied as the molecule is a target for antibiotic development. There are four possible types of PG cleavage: N-Acetyl-β-glucosaminidases (cleaves the glycosidic bond between NAG-NAM), muramidase (lysozyme which is a hydrolase that produces a reducing NAM product, and lytic transglycosylase that produces anhydro-acetylmuramic acid), peptidase (carboxypeptidase and endopeptidase), and amidase (hydrolyzes the amide bond between NAM and the oligopeptide) [8–10].

Canonically, endolysin molecules consist of an N-terminal enzymatic catalytic domain (ECD, i.e., amidase and peptidase) and a C-terminal cell wall binding domain (CBD, such as LysM and PG-bd-like) [11–13]. In the late stage of a phage cycle, endolysin accumulates in the cytosol while phage assembly takes place. The accumulated endolysin molecules eventually leave the cytosol and degrade PG as a result of the membrane pore formation initiated by aggregation of holin molecules [14]. However, it has been reported that some coliphage endolysins have the Signal-Anchor-Release (SAR) domain. It is a weak transmembrane region located in the N-terminus of the protein. The first discovered SAR endolysin, coliphage P1 Lyz, was shown to be cysteine dependent [15]. When the protein is synthesized, this domain signals the protein to co-localize with the inner membrane of the host where it is arrested and inactivated. As P1 pinholin disrupts the membrane proton motive force, the SAR endolysin leaks out from the cell membrane. Ultimately, disulfide bonds are formed leading to active site formation [1, 14–16].

To date, most of the well-characterized endolysins are those from enterobacteriophages such as coliphage T4, T7, P1, and *Salmonella* phage SPN1S [15, 17–19]. Other well-characterized endolysins include those from *Streptomyces aureofaciens* phage μ1/6, *Streptomyces avermitilis* phage phiSASD1, *Streptococcus pneumoniae* phage Cpl-7, *Burkholderia cenocepacia* phage Ap3, and mycobacteriophage D29 [20–25].

Through the SEA-PHAGES program [26], more than 300 genomes of *Streptomyces* phages have been added to the Actinobacteriophage Database [27], but few of the genes have been experimentally characterized [21, 23]. Here, we utilized bioinformatic analyses to identify potential proteins of interest. The functional domains of 250 annotated endolysin protein sequences from these *Streptomyces* phages were predicted and classified into nine different combinations of ECD and CBD. Three proteins of interest with different predicted enzymatic activities were subsequently selected for gene cloning and protein production. We hypothesized that (1) the proteins of interest will have the ability to degrade purified PG, and (2) the proteins will be potential antimicrobial agents against ESKAPE safe relatives.

## 2 Results and Discussion

### 2.1 Bioinformatic analysis

Bioinformatic analysis was used to identify the diversity of endolysins within phages that infect *Streptomyces*. Results from domain prediction have shown that 250 protein sequences of the putative endolysins/lytic enzymes used in this study can be classified into nine different types based on combination of the ECD and CBD (Figure 1A). Predicted enzymatic functions include amidase (PGRP, amidase, and CHAP), peptidase (peptidase and CHAP), and transglycosylase (GH108)/glycosyl hydrolase (Figure 1B). These results were used to select three putative endolysins of interest for experimental studies due to the differences in their catalytic and binding domains, including LazerLemon gp35 (CHAP and PG-binding), Nabi gp26 (amidase and LysM), and Tribute gp42 (PGRP/amidase and LysM); all of which are from phages isolated on the host bacterium *Streptomyces griseus* ATCC 10137.

**FIGURE 1.**
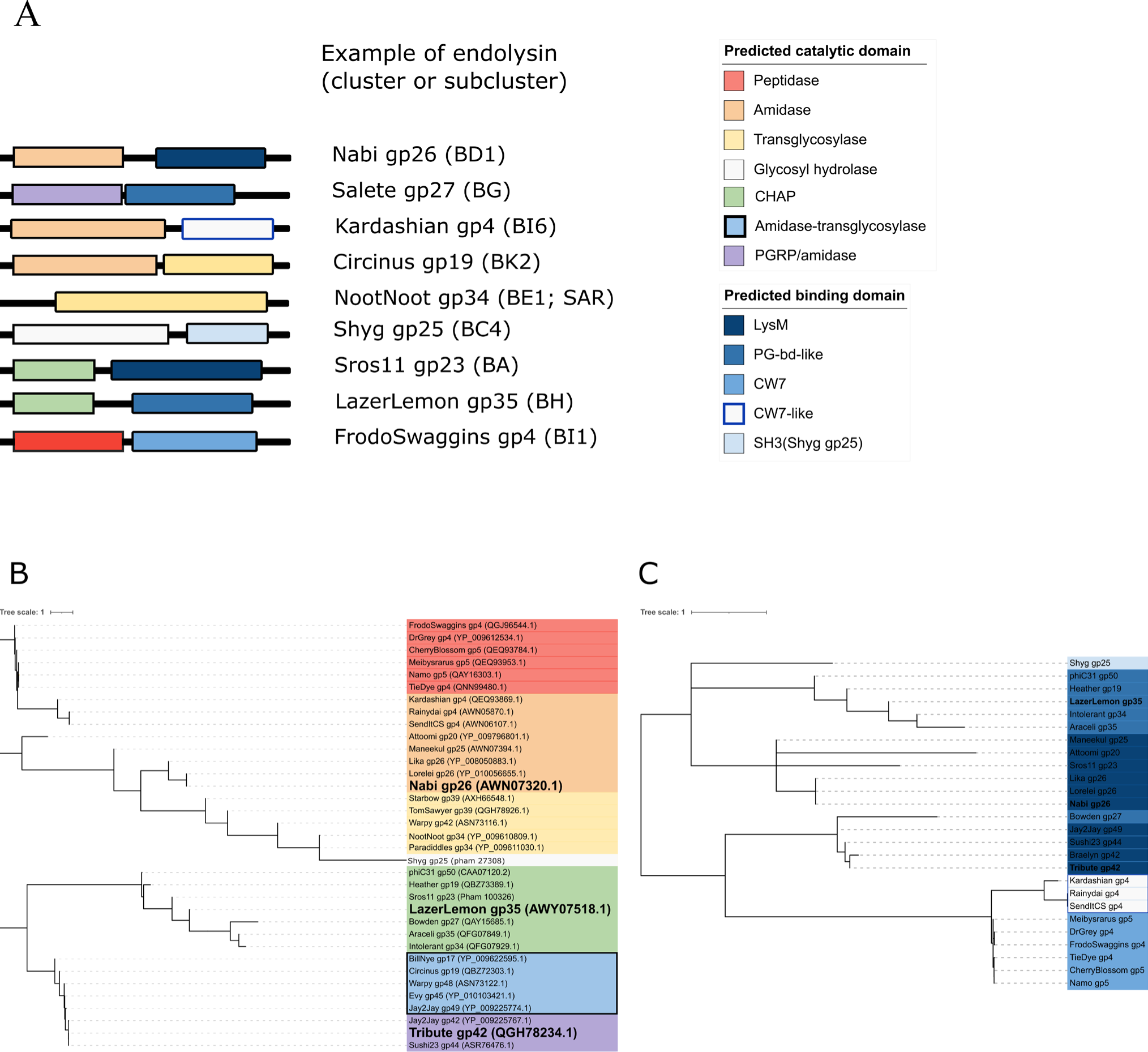
Domain prediction and phylogeny. **(A)** Domain modular structures of the putative endolysins show catalytic domain at the N-termini while binding domain is at the C-termini (except Circinus gp19). A total of nine different types were found based on the combination of ECD and CBD. **(B)** Phylogenetic trees generated by ECD and CBD alignments. Boldface fonts indicate proteins of interest.

### 2.2 Protein sequence information

The 250 putative endolysins are small proteins with the approximate molecular mass of 18.8 – 50.8 kDa (Additional file 1). It was previously reported that the shortest *Streptomyces* phage putative endolysin had 226 aa residues [28]. However, the current study utilized a larger dataset and found 31 small putative endolysins (174 -197 residues) representing second endolysins/lytic proteins that might have been previously overlooked in the cluster BE genomes (Figure 2A). Interestingly, results obtained from the Center for Phage Technology (CPT) Galaxy [29] have suggested that these endolysins (n=31) possess a SAR-like domain at the N-terminus (Figure 2B). Of these putative SAR-like proteins in this study, seven are possibly cysteine-independent (Additional file 2). To date, putative SAR endolysins from Gram-positive systems are rare. Only four from lactococcal phages have been previously reported [30].

**FIGURE 2.**
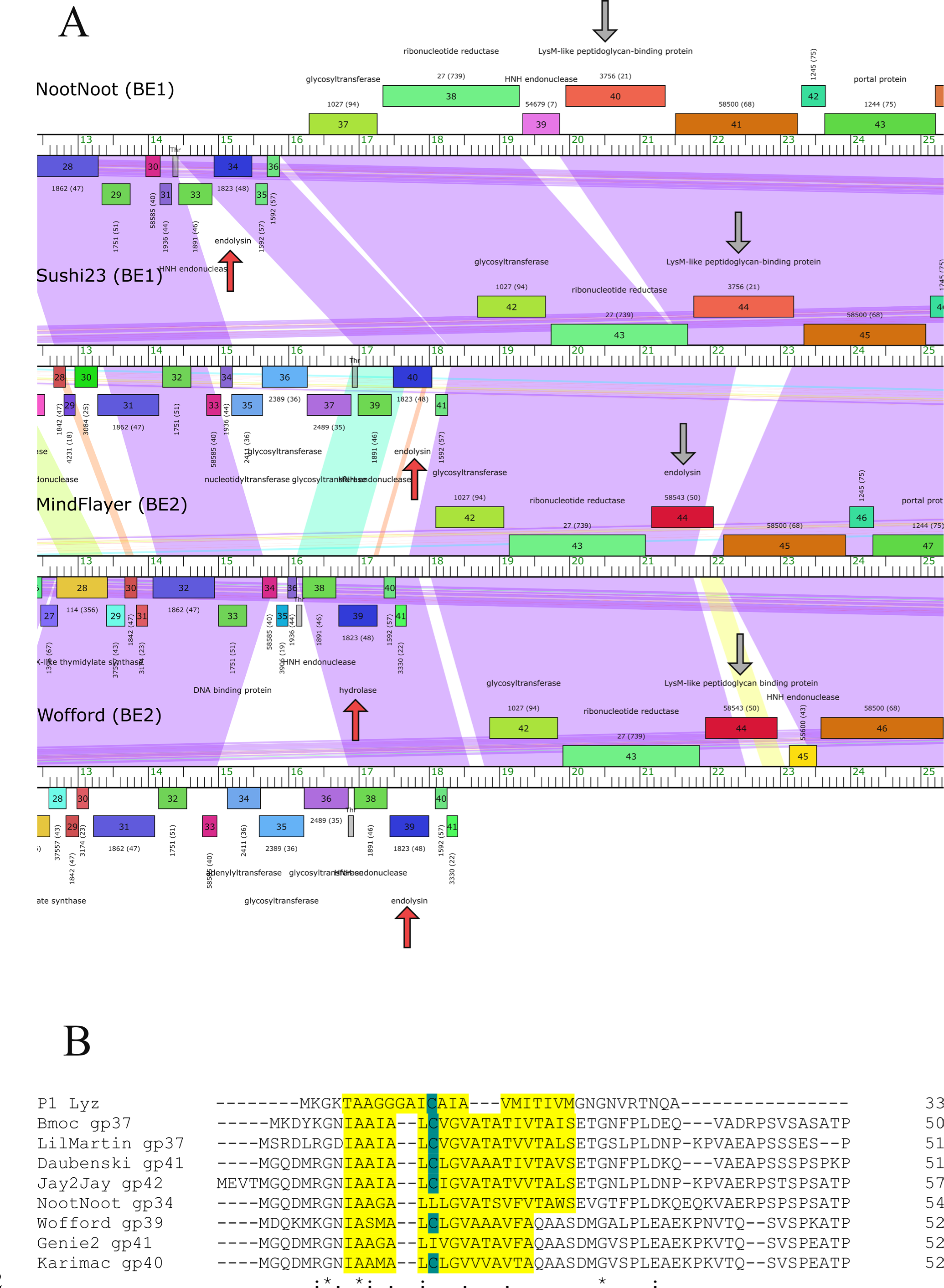
The small putative endolysins on cluster BE genome map were predicted SAR-like endolysins. **(A)** Partial genome map alignment of cluster BE phages shows two different endolysin gene locations (Indicated by arrow. Red arrows = predicted SAR). Rulers represent the length of the genome (kbp). Phage names are shown on the left top of each genome. Data was visualized on Phamerator [41]. **(B)** Protein sequence alignment of P1 Lyz and our predicted SAR endolysins. Putative SAR domain is highlighted in yellow and cysteine residues are shown in blue.

### 2.3 Molecular cloning and protein production

Each gene of interest was inserted between the T7 promoter and terminator of each pET-28a(+) construct and was confirmed by colony PCR and Sanger sequencing. Proteins expression was induced from three different clones (*E. coli* BL21(DE3)) each containing a corresponding insert. Results from purification and SDS-PAGE show successful production of three recombinant enzymes, including gene products from LazerLemon gp35 (LL35lys), Nabi gp26 (Nb26lys), and Tribute gp42 (Tb42lys). The level of protein expression and purity was also assessed by SDS-PAGE (Figure 3). Interestingly, LL35lys showed three purified products including the full-length protein (31 kDa), and two shorter fragments (24 and 22 kDa, approximately). It has been reported that CHAP endolysins from clostridial and staphylococcal phages can produce truncated CBD assisting in enzymatic activity [31, 32]. However, our recombinant enzymes contain a 6X His tag at the N-terminus. Thus, the purified fragments should not be truncated CBDs. The estimated protein sizes of these two have suggested that they potentially include the entire ECD.

**FIGURE 3.**
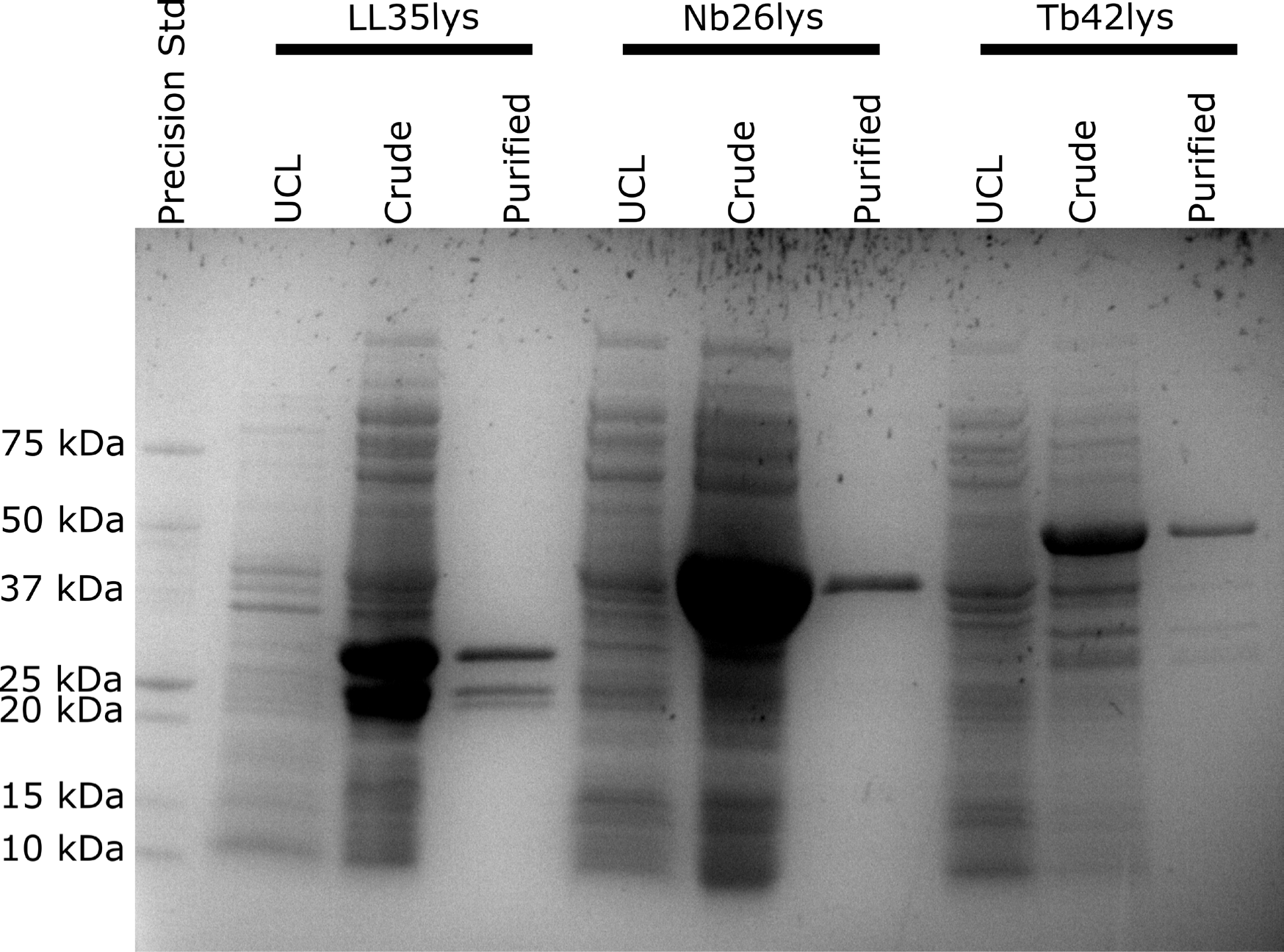
SDS-PAGE result. Recombinant protein expression and purity were evaluated by SDS-PAGE. When compared to uninduced sample (UCL), proteins were overexpressed after 2-4 h of IPTG induction. High purity was observed (except some contaminations in the Tb42lys lane). LL35lys lane consisted of three different products: 31 kDa (full length), 24-, and 22-kDa segments. Expected band sizes are 31.1 kDa (6xHis-T7-LL35lys), 42.6 kDa (6xHis-T7-Nb26lys), and 54.5 kDa (6xHis-T7-Tb42lys).

Using this protein expression system and inducing at 30°C for 2-4 h, LL35lys and Nb26lys yielded approximately 14-16 mg per one litter of culture. However, Tb42lys has a significant lower yield (≤1 mg/1L of culture). When the concentration of imidazole used for elution was increased up to 1 M, the yield for Tb42lys was slightly improved (up to 2 mg/1L of culture).

### 2.4 PG-degrading activities

To test PG degradation activity of the recombinant proteins, change in PG turbidity was analyzed using two different methods: zymography and hydrolysis assay. Zymography was carried out using gel electrophoresis and visualization of bands where enzymatic degradation occurred. To provide the substrate, micrococcal PG was mixed in the acrylamide gel solution prior to polymerization, creating a homogenously turbid gel. After electrophoresis and washes with dH_2_O to remove SDS/buffer, especially Tris ((HOCH_2_)_3_CNH_2_) which was previously reported to inhibit activity of carbohydrate hydrolases, aminopeptidases, aminotransferases, and cholinesterases [33], LL35lys (including 31-, 24-, and 22-kDa fragments), Nb26lys, and Tb42lys were able to create clear zones at the corresponding protein migration distance (Figure 4), indicating PG-degradation activities. The presence of clearing in the zymogram for the 24-, and 22-kDa segments from LL35lys suggests that these are not truncated CBDs but include the full ECDs. However, the relevance of these bands to in vitro activity is unknown and would require further investigation.

**FIGURE 4.**
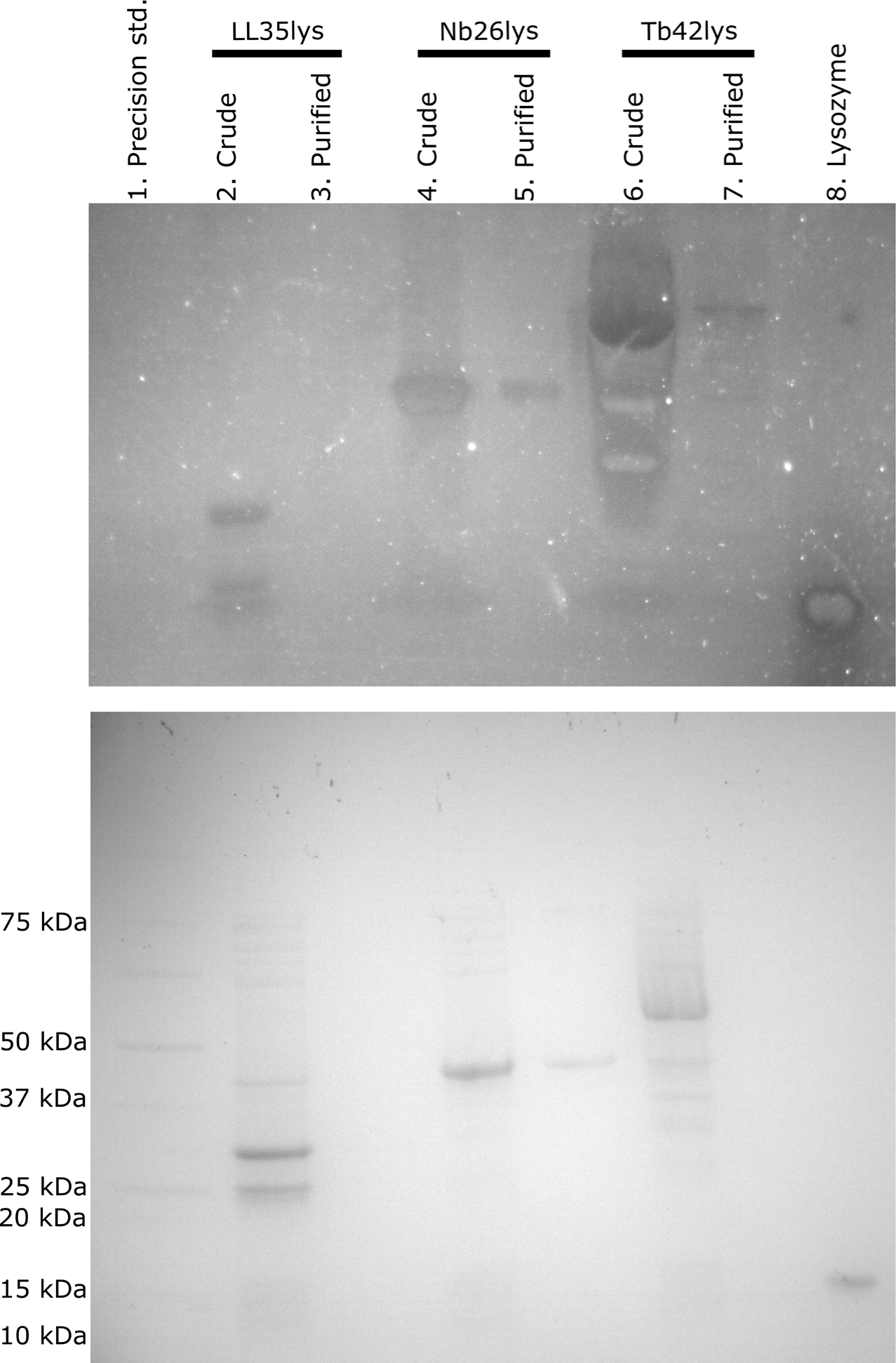
Zymography analysis. Without gel staining (top panel), the enzymes were able to degrade micrococcal PG embedded in acrylamide gel (10% hand cast), illustrating as clear bands (lane 2, and 4-8). In contrast, while crude Tb42lys (lane 6) showed false positives due to high sensitivity/concentration, LL35lys was easily precipitated and showed negative results (lane 3). In the lower panel, the proteins were separated by SDS-PAGE (4-20% Mini-PROTEAN® TGX™) and stained with Coomassie blue, serving as an annotated gel. The Precision standard (lane 1) was used as a negative control for zymogram, while commercial lysozyme (lane 8) serves as positive control. Lanes 2, 4, and 6 represent crude extracts from *E. coli* (BL21(DE3)) induced cultures. Lanes 3, 5, and 7 are purified proteins.

The hydrolysis assay results demonstrated that the recombinant enzymes can degrade micrococcal PG at a much slower rate than commercial lysozyme (Figure 5). At room temperature, lysozyme can completely hydrolyze the substrate as soon as five minutes (Figure 5B), while our proteins take at least two hours to start the PG degradation (turbidity reduction to 32-40%; Figure 5A and 5C). This is consistent with studies in endolysins from mycobacteriophage (amidase) and enterobacterial phage (lysozyme) which were reported to take 1-2 h to degrade PG [135, 136]. Note that the hydrolysis assay performed in this study was carried out under the same conditions (PBS pH 7.4, 37°C) for all proteins. ZnCl_2_ with a final concentration of 2 mM was supplemented in another reaction of PG-Nb26lys. However, the metal ion appeared to interfere with the reaction leaving more turbidity (50%) remaining after the treatment (Figure 6A). This indicates that Nb26lys might not be zinc amidase as suggested by the bioinformatic results (Figure 6B).

**FIGURE 5.**
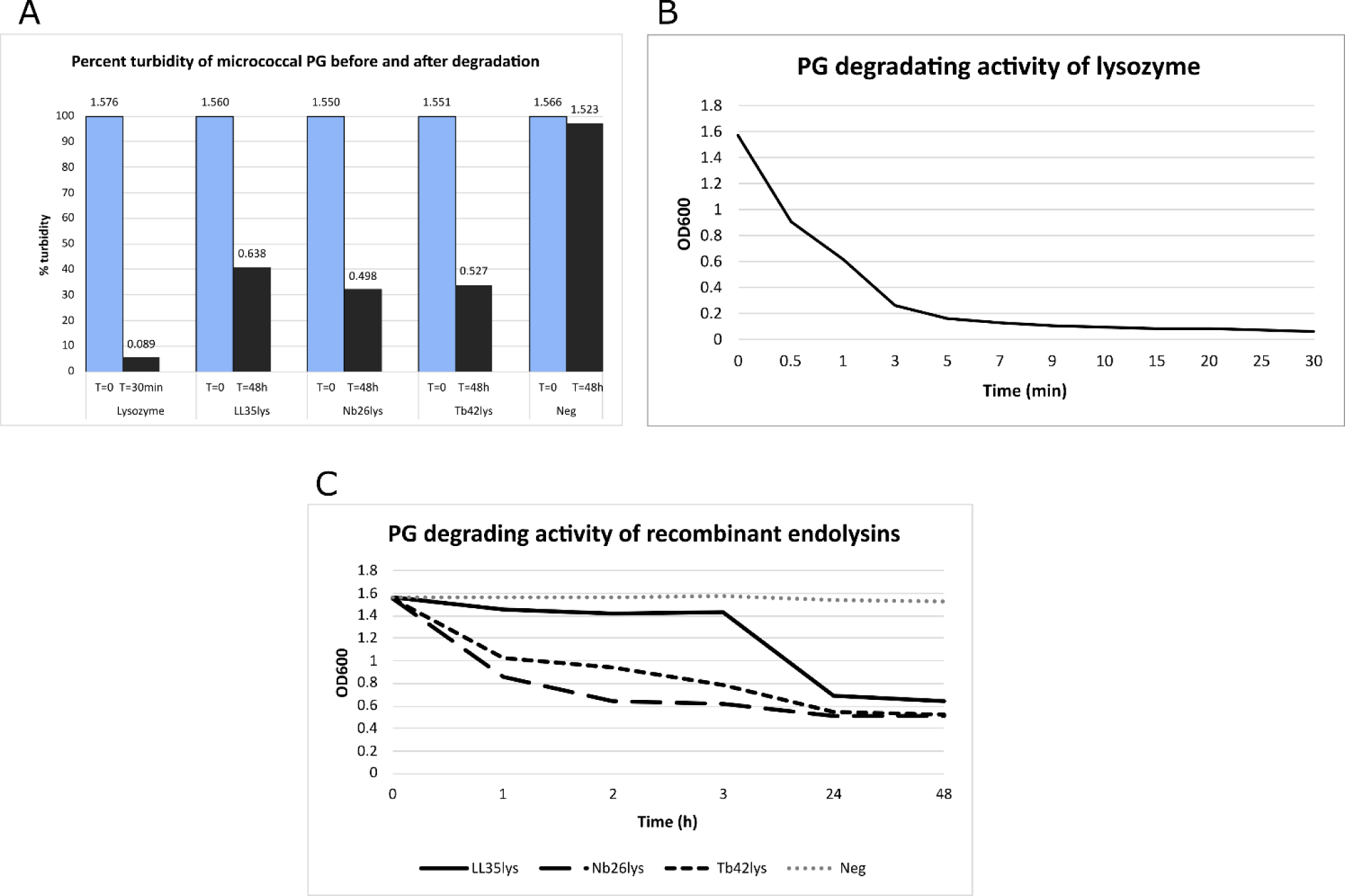
Hydrolysis results. The micrococcal PG was hydrolyzed in PBS (pH 7.4 at 37°C) by different recombinant endolysins. **(A)** Percent turbidity shows that lysozyme can reduce the insoluble PG to 5%, while the recombinant endolysins can reduce the turbidity to approximately 32-40%. The figure above each bar column indicates OD_600_ results. Blue and grey columns show results before and after the enzyme reaction. **(B)** Hydrolysis reaction of lysozyme **(C)** Hydrolysis reaction of the recombinant enzymes shows that Nb26lys and Tb42lys degraded PG at a faster rate than that of LL35lys.

**FIGURE 6.**
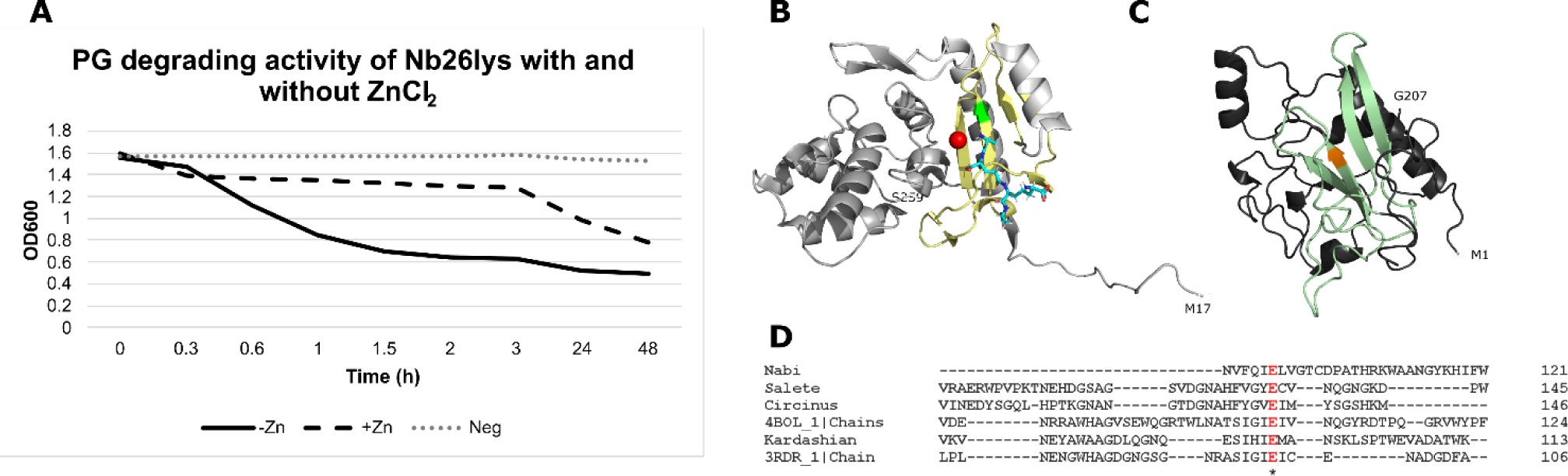
ZnCl_2_ interfered with PG degradation of Nb26lys. **(A)** Hydrolysis assay showed that at a low concentration of 2 mM, ZnCl_2_ slowed down the reaction approximately ten-fold (from 0.3 to 3 h). **(B)** Structure of AmpDh2, a zinc protease that plays a role in PG turnover from *Pseudomonas aeruginosa* (PDB 4BOL). Glu106 (green) has been reported to be a conserved residue at the active site (five yellow sheets) [11]. Zinc ion is illustrated as a red sphere and the substrate is a blue stick. **(C)** AlphaFold predicted structure of Nb42lys ECD showing potential active site (four sheets; light green) that is aligned with that of AmpDh2. Glu98 is labeled orange. **(D)** Glu98 of Nb26lys has shown to be aligned with Glu106 of AmpDh2. Nb42lys ECD and 4BOL chain A are 24% identical.

### 2.5 Antimicrobial activities

#### 2.5.1 Killing assay

Based on the three recombinant proteins’ ability to degrade micrococcal PG, we hypothesized that the proteins would be potential antimicrobial agents against *S. griseus* and ESKAPE safe relatives. To screen for antimicrobial activity, we employed the killing assay. The cells were treated with three different concentrations (100, 200, and 400 µg/mL) of each enzyme. PBS and cells were used as buffer control. Crude extracts from uninduced samples were also tested for negative control. At each time point of exposure, the sample was dropped onto Luria agar (LA) and subsequently incubated at 30°C overnight to examine bacterial growth. As expected, the three enzymes were able to kill the phage isolation host, *S. griseus*. Interestingly, at a lower concentration of 100 µg/mL, the growth level was increased when treated for 24 h. This might indicate that the proteins could be recycled by the *Streptomyces* cells at sub-lethal concentrations. Importantly, this phenomenon does not occur with the ESKAPE safe strains.

In our study, the killing assay has revealed that, while Nb26lys and Tb42lys are originally from a Gram-positive system, they show killing of Gram-negative strains, with Tb42lys effective against three strains (*E. coli*, *P. putida, A. baylyi* and *K. aerogenes*) (Table 1 and Additional file 3). Within 2 h of exposure, Tb42lys can completely kill *E. coli* and *P. putida*. Nb26lys can also kill *E. coli* and *K. aerogenes* within 2 and 24 h, respectively. Furthermore, Tb42lys requires longer than 2 h to kill *A. baylyi* and *K. aerogenes.* It is important to note that commercial lysozyme is not effective in killing these strains without any other chemical or mechanical approaches. This result suggests that Tb42lys and Nb26lys may have a greater ability to penetrate the outer membrane and access the PG layer. It has been reported that antibiotics, organic acid, SAR domain, and nanoparticles are some examples of technologies that have been reported to improve the permeability [34].

**Table 1.** Lowest growth level of *S. griseus* and ESKAPE after endolysin treatment.

| Strains | LL35lys |  |  | Nb26lys |  |  | Tb42lys |  |  | Lysozyme |  |  |
| --- | --- | --- | --- | --- | --- | --- | --- | --- | --- | --- | --- | --- |
|  | Activity | Duration | LGL | Activity | Duration | LGL | Activity | Duration | LGL | Activity | Duration | LGL |
| Sg | Y | 1h | + | Y | 1h | - | Y | 2h | - | Y | 10min | - |
| E | Y | 24h | + | Y | 24h | + | Y | 24h | ++ | Y | 24h | - |
| S | Y | 24h | + | Y | 24h | ++ | Y | 24h | ++ | N | N/A | +++ |
| K | N | N/A | +++ | Y | 24h | - | Y | 24h | - | N | N/A | +++ |
| A | N | N/A | +++ | Y | 24h | ++ | Y | 24h | - | N | N/A | +++ |
| P | N | N/A | +++ | N | N/A | +++ | Y | 2h | - | N | N/A | +++ |
| E | N | N/A | +++ | Y | 2h | - | Y | 2h | - | N | N/A | +++ |

Sg; *S. griseus*, E; *Enterococcus raffinosus*, S; *Staphylococcus epidermidis*, K; *Klebsiella aerogenes*, A; *Acinetobacter baylyi*, P; *Pseudomonas putida*, and E; *Escherichia coli*. LGL; lowest growth level

#### 2.5.2 Minimal inhibitory concentration (MIC)

To quantify potential antimicrobial activity of our recombinant endolysins, MICs were determined. Due to the low yield of Tb42lys, only LL35lys and NB26lys were measured. MIC are shown for LL35lys treatment of *E. raffinosus* and *S. epidermidis* and Nb26lys treatment of *E. coli* and *K. aerogenes* (Figure 7). Other enzyme/bacteria combinations did not show similar reductions. Within the range of 2 to 300 µg/mL endolysin concentration, 300 µg/mL of Nb26lys showed inhibition of approximately 80% and 99% growth of *E. coli* and *K. aerogenes*, respectively. Concentrations below 150 µg/mL did not reach a 50% reduction in growth. In contrast, LL35lys at 2 µg/mL was able to inhibit approximately 90% of *S. epidermidis* and *E. raffinosus* growth. Of the endolysin treatments currently under consideration by U.S. Food and Drug Administration, Endolysin SAL-1 (formulated as SAL-200 or Tonabacase) has shown lower MICs of 0.22-1.24 µg/mL against several strains of *S. aureus*, while LysK (CHAP-amidase-SH3b cell wall binding domain), an endolysin that is very similar to SAL-1 showed the MICs of 0.44-6.52 µg/mL [35, 36]. Previous studies have demonstrated that the presence of 500 µg/mL of endolysins from *A. baumannii* phages can reduce pathogen growth to as low as 1% (equivalent to MIC_99_) [37, 38]. Altogether, these results indicate the promise that *Streptomyces* phage endolysins LL35lys and Nb26lys could be effective as an antimicrobial agent and should receive further study.

**FIGURE 7.**
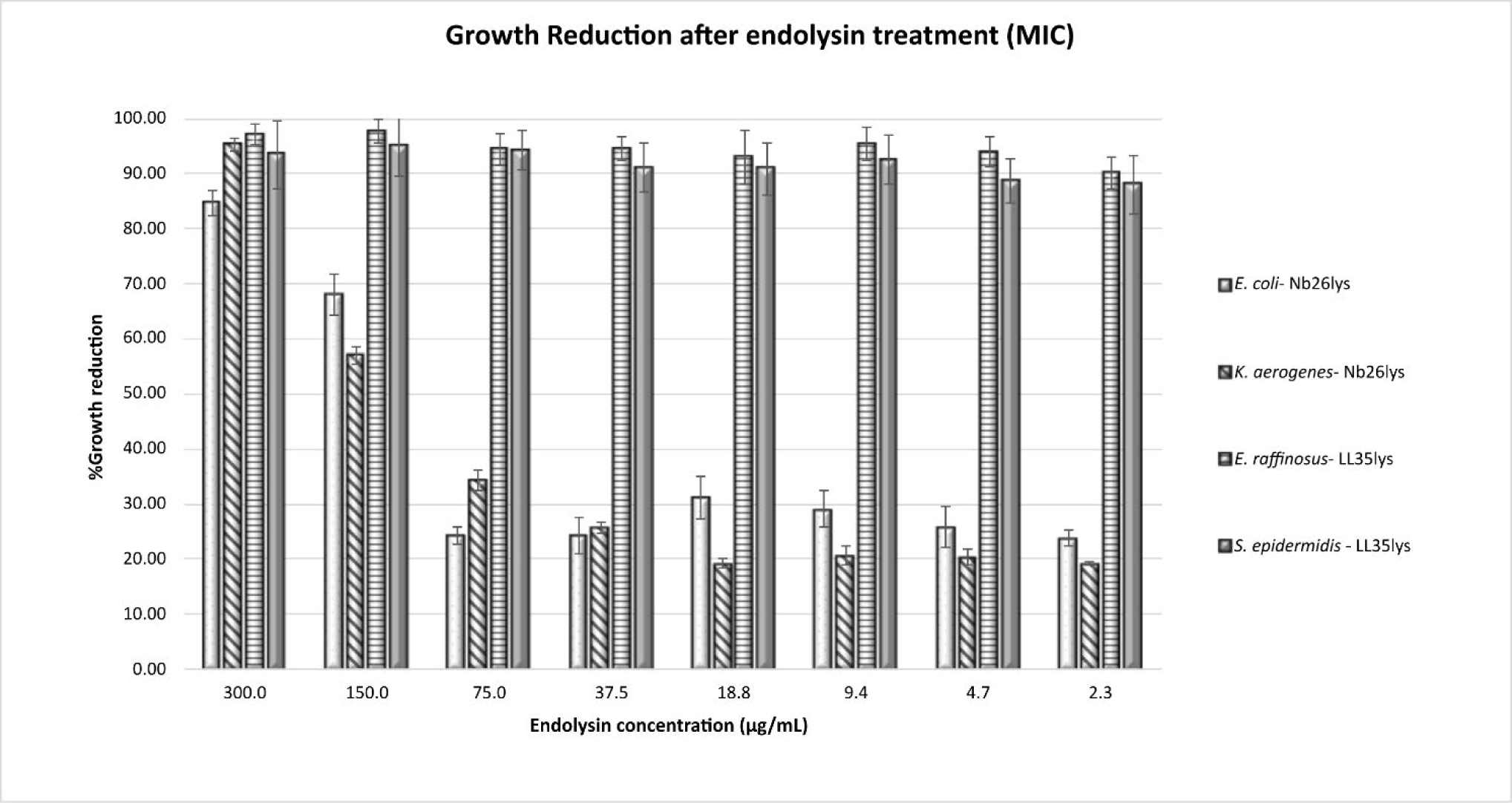
MIC results. Within a range of 2 to 300 µg/mL of LL35lys and Nb26lys, the results showed that 2 µg/mL of LL35lys can inhibit approximately 90% of *E. raffinosus* and *S. epidermidis.* However, a higher concentration of 300 µg/mL of Nb26lys is required to treat *E. coli* (MIC_85_) and *K. aerogenes* (MIC_95_).

## 3 Conclusion

Our study demonstrates the potential to use bioinformatic tools to prospect biomolecules from phages that infect non-pathogenic microorganisms such as *Streptomyces*. 250 putative endolysins from *Streptomyces* phages were classified into 9 different types based on the functional domain combination. Endolysins from the phages LazerLemon, Nabi, and Tribute were selected for experimental studies. LL35lys and Nb25lys were successfully extracted and purified with the approximate yield of 14-17 mg per liter of culture, while only 1-2 mg per liter of culture was yielded from Tb42lys. All three recombinant enzymes showed micrococcal-PG degradation on zymogram. The hydrolysis results suggest that 100 µg/mL of enzymes can reduce the PG turbidity to 32-40% within 48 h. Furthermore, Tris, the widely used buffer, has been shown to interfere with the enzyme activity, especially that of LL35lys. Additionally, the hydrolysis experiment revealed that 2 mM of ZnCl_2_ temporarily suppressed the activity of Nb26lys. Results from killing assays have suggested that other than bactericidal effect on *S. griseus*, Nb26lys and Tb42lys can reduce the growth of Gram-negative ESKAPE relatives, while LL35lys are more specific to the Gram-positive strains with a low MIC_90_ of 2 µg/mL. A concentration of ≥300 µg/mL of Nb26lys is required to inhibit *P. putida* and *K. aerogenes*. Thus, endolysins from *Streptomyces* phage have competence as potential antimicrobial agents against *S. griseus* and ESKAPE safe relatives.

## 4 Methods

### 4.1 Bioinformatics

The 250 putative endolysin protein sequences were previously annotated by SEA-PHAGES [26] participants, and available on Phamerator [42] as collected from the Actinobacteriophage Database [27]. The list of the Genbank accession numbers for the genome sequences can be found in Additional file 1. Functional domains were predicted by InterPro v83.0 [43] and HHpred^1^ [39] with default settings. Predicted 3D structures were generated from Alphafold2 Colab v2.3.2 [44]. SAR Finder developed by the CPT Galaxy [29] was used to search for SAR domains. As well, molecular mass, length, isoelectric point, and amino acid composition of the protein sequences were analyzed using Expasy ProtParam [45], IPC 2.0 [40], and Antheprot software v6.9.3 [46]. To generate phylogenetic trees, protein sequences covering ECDs or CBDs were aligned using ClustalW. The trees were constructed using Neighbor-Joining Tree Test with a bootstrap of 500 by MEGA 11 software v11.0.13 [47].

### 4.2 Phage propagation and bacterial growth conditions

Phages LazerLemon (Genbank accession number: MH229865.1), Nabi (MH171094.1), and Tribute (MN369743.1) were previously isolated from soil samples and sequenced by SEA-PHAGES participants using *Streptomyces griseus* ATCC 10137 as the isolation host. Information on phage discovery can be found from the Actinobacteriophage Database [27]. A small amount (10-20 µL) of the selected phage lysates was withdrawn from our phage collection at the University of North Texas. Phage propagation and DNA extraction were performed according to standard protocols [48]. *E. coli* BL21(DE3) was used as an expression system (Novagen #70777) and maintained according to the manufacturer’s instructions. ESKAPE safe relatives used in this study consists of *Enterococcus raffinosus* ATCC 49464, *Staphylococcus epidermidis* ATCC 12228, *Klebsiella aerogenes* ATCC 13048, *Acinetobacter baylyi* ATCC 33304, *Pseudomonas putida* ATCC 12633, and *Escherichia coli* ATCC 25922. All ESKAPE strains were grown at 30-37°C using Luria broth/agar (BD Difco, Fisher Scientific #DF0145-17-0) for 15-24 h.

### 4.3 Cloning, protein expression, extraction, purification, and storage

Primers used for PCR can be found in Table 2. The amplified and digested inserts were ligated into pET-28a(+) (Novagen #70777) before transformation into chemically competent BL21(DE3) by heat shock. To maintain the selected clones, kanamycin (50 µg/mL) was supplemented in the medium as a measure of antibiotic selection. All clones were confirmed by colony PCR and Sanger sequencing. In a one-litter culture (OD_600_ of 0.4-0.6), protein expression was induced by 0.5 mM IPTG at 30°C for 3-4 h. After centrifugation at 4,000 x g, 4°C for 30 minutes, the cell pellet was resuspended in Tris buffer (20mM Tris and 250 mM NaCl, pH 8.0) supplemented with Halt^TM^ protease inhibitor (ThermoFisher #87786). The cells were subsequently lysed using freeze-thaw cycles. Cell lysate was then extracted by centrifugation at 21,000 x g for 30 minutes and purified using HisPur Ni-NTA columns (Thermo Scientific #88224). The bound proteins were eluted from the column using 250-750 mM imidazole elution buffer (20 mM sodium phosphate and 300 mM NaCl, pH 7.4). Spectrophotometry and SDS-PAGE were used to determine the protein concentration and purification. Small aliquots of protein samples were fast frozen in liquid nitrogen before storing at - 80°C.

**Table 2.**
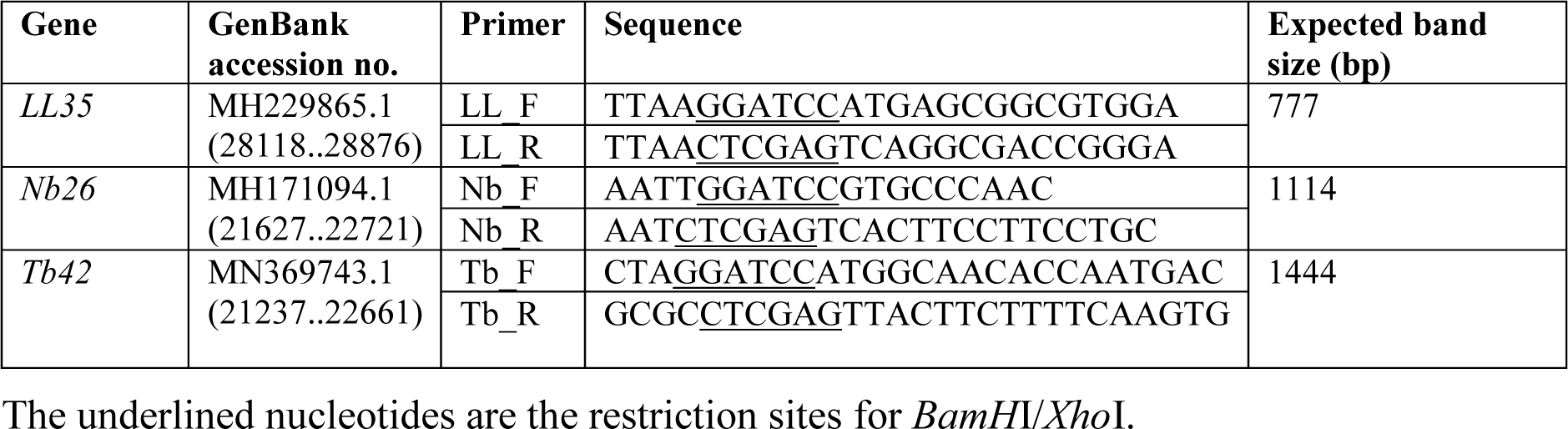
Specific primers used for gene amplification.

### 4.4 Zymography

Zymography in this study was carried out with modifications from previous studies [3, 48]. A 10% acrylamide gel was hand-cast without addition of SDS. Prior to polymerization, 5 mg/mL of PG from *Micrococcus lysodeikticus* ATCC No. 4698 (Sigma Aldrich #M3770) was added directly to the gel solution. Approximately 2-5 µg of the non-denatured protein samples were diluted 1:1 with the native sample buffer (BIO-RAD #161-0738) and loaded into the polymerized gel. Tris/Glycine/SDS buffer (BIO-RAD #161-0732) was used as a running buffer. The gel was electrophoresed at 100V for approximately 1 h, before washing in DI water to remove SDS and buffer components. Because Tris can interfere with LL35lys and Nb26lys activities, the gel was soaked in fresh DI water at 30°C. After 2-3 h of incubation, a change of water was done to allow removal of any remaining Tris residuals. The gel continued soaking in the fresh DI water overnight at 30°C. Without staining, the zymogram gel was visualized in a dark room using white light and a dark background.

### 4.5 Hydrolysis assay

The hydrolysis reaction was set up in a total volume of 1 mL. Micrococcal PG was dissolved in pH 7.4 PBS to a final concentration of 1 mg/mL and pre-warmed at 37°C. Using the BioSpec-mini spectrophotometry, OD_600_ was recorded as a T=0 reading before adding 100 µg/mL of the enzyme. After adding the enzyme to the pre-warmed PG solution, it was incubated at 37°C for 24 h. OD_600_ was recorded at intervals of 10 seconds for lysozyme, and 30 minutes for the recombinant enzymes.

### 4.6 Killing assay

In 50 µL treatments, different concentrations of each recombinant protein were added to molecular-grade water, followed by 3 µL of the fresh overnight culture. Buffer control consisting of cells and water was tested parallel with the treatment. Prior to and after incubation of the liquid mixture at 30°C, 3 µL of sample was dropped onto LA which was subsequently incubated at 30°C overnight to grow any surviving cells. The different levels of cell growth observed on the plate were recorded as “+++” for highest level, “++” for moderate level, “+” for low level, and “-” for no growth.

### 4.7 Minimal inhibitory concentration (MIC)

MIC in this study was performed using a microdilution method in liquid medium as previously described [49, 50]. Approximately 10^5^ cfu/mL of fresh overnight cultures of *S. griseus* and ESKAPE relatives were treated with the recombinant endolysins which were serially diluted by 2-fold in a 96-well microplate. Blank control was carried out by adding enzyme to PBS, while growth control was a mixture of cells and the same buffer. The treatments were subsequently incubated at 30°C, overnight. A microplate reader (BioTek Synergy Mx SMA, Fisher Scientific #FISBTSMATD) and Gen5 software v2.01 were used to obtain OD_600_ data.

## Supporting information

Additional file 1 List of the 250 putative endolysins

## Supplementary Information

**Additional file 1** List of the 250 putative endolysins used in this study.

**Additional file 2** Putative topological and SAR domain found in the predicted SAR endolysins form phages cluster BE.

**Additional file 3** Growth level of ESKAPE and *S. griseus* after endolysin treatment (killing assay).

## 5 Declarations

### 5.1 Ethics approval and consent to participate

Not applicable

### 5.2 Consent for publication

Not applicable

### 5.3 Availability of data and materials

The phage genomes supporting the conclusion of this article is available in GenBank, [https://www.ncbi.nlm.nih.gov/genbank/].

### 5.6 Authors’ contributions

JM and LH wrote and edited the main manuscript text. AC did quality control of the bioinformatic data. RH, FL, and JS assisted in molecular cloning and protein expression. Other authors gathered protein sequences from the database and were involved in bioinformatic analyses. JM carried out bioinformatic observations, experimental studies, discussions, and graphic images. All authors reviewed the manuscript.

## 5.7 Acknowledgments

The authors would like to thank the SEA-PHAGES program for the resources from Phamerator and PhagesDB, and Toulouse Graduate School, University of North Texas for the Graduate Student Research Award.

---

**Additional file 1** List of the 250 putative endolysins with protein information and GenBank accession number.

**Additional file 2** Putative topological and SAR domain found in the predicted SAR endolysins form phages cluster BE. The results were generated by CPT Galaxy showing 31 predicted SAR endolysins reported in this study.

**Additional file 3** Growth level of ESKAPE and *S. griseus* after endolysin treatment (killing assay). Bacterial cell cultures were treated with a different concentration of LL35lys, Nb26lys, and Tb42lys. Samples were dropped onto agar after 10 min, 1 h, 2h, and 24 h of the treatment. Reduction in colonies shows positive result, when compared to the control.

## Footnotes

1 Version 57c8707149031cc9f8edceba362c71a3762bdbf8

