## Additional file 1 List of the 250 putative endolysins for "Investigating Novel *Streptomyces* Bacteriophage Endolysins as Potential Antimicrobial Agents"

Table S1

**Table S1.** 250 putative endolysins analyzed in this study.

| No | Endolysins | Genbank/pham Accession no. | Phage Cluster | Subcluster | AA length | Mass (kDa) | Predicted Function (by SEA-PHAGES) | Notes |
|---|---|---|---|---|---|---|---|---|
| **Amidase-LysM** | | | **20.0%** | | | | | |
| 1 | >SPB78 gp21 | NZ_ACEU00000000.1 | BA | None | 275 | 29.0 | N/A | |
| 2 | >Nabi gp26 | AWN07320.1 | BD | BD1 | 364 | 39.0 | LysM-like endolysin | |
| 3 | >Toma gp26 | AWN07621.1 | BD | BD1 | 362 | 38.7 | LysM-like endolsyin | |
| 4 | >Asten gp25 | QAY17707.1 | BD | BD1 | 362 | 38.5 | LysM-like endolysin | |
| 5 | >Goby gp26 | AWN07545.1 | BD | BD1 | 362 | 38.7 | LysM-like endolysin | |
| 6 | >Lika gp26 | AGM12049.1 | BD | BD1 | 364 | 39.1 | N/A | |
| 7 | >Whatever gp25 | QFP95192.1 | BD | BD1 | 362 | 38.5 | lysin A | |
| 8 | >Godpower gp27 | AOQ27003.1 | BD | BD1 | 364 | 39.1 | endolysin | |
| 9 | >Dattran gp27 | ATE85131.1 | BD | BD1 | 362 | 38.7 | endolysin | |
| 10 | >Lorelei gp26 | AOQ26925.1 | BD | BD1 | 364 | 39.0 | endolysin | |
| 11 | >Danzina gp27 | YP_009592392.1 | BD | BD1 | 362 | 38.7 | LysM | |
| 12 | >Brataylor gp28 | AOQ27076.1 | BD | BD1 | 364 | 38.9 | endolysin | |
| 13 | >Zemlya gp 27 | AGM12202.1 | BD | BD1 | 362 | 38.7 | N/A | |
| 14 | >Sujidade gp27 | AGM12125.1 | BD | BD1 | 362 | 38.8 | N/A | |
| 15 | >Yasdnil gp25 | YP_010056432.1 | BD | BD1 | 362 | 38.5 | LysM-like endolysin | |
| 16 | >TuanPN gp25 | QDK03196.1 | BD | BD1 | 362 | 38.5 | LysM-like endolysin | |
| 17 | >OzzyJ gp26 | AVE00407.1 | BD | BD1 | 362 | 38.5 | LysM-like endolysin | |
| 18 | >Rana gp26 | AWN07244.1 | BD | BD1 | 364 | 39.0 | LysM-like endolysin | |
| 19 | >Maneekul gp25 | AWN07394.1 | BD | BD1 | 362 | 38.5 | LysM-like endolysin | |

*(table continues)*

| No | Endolysins | Genbank/pham Accession no. | Phage Cluster | Subcluster | AA length | Mass (kDa) | Predicted Function (by SEA-PHAGES) | Notes |
|---|---|---|---|---|---|---|---|---|
| **Amidase-LysM** | | | **20.0%** | | | | | |
| 20 | >Celeste gp26 | ATE85054.1 | BD | BD1 | 362 | 38.7 | endolysin | |
| 21 | >Teutsch gp44 | QAX95780.1 | BE | BE1 | 474 | 50.8 | LysM-like PGBD | |
| 22 | >Mildred21 gp42 | YP_009610583.1 | BE | BE1 | 336 | 36.4 | LysM-like PGBD | |
| 24 | >Paradiddles gp40 | YP_009611036.1 | BE | BE1 | 471 | 50.4 | LysM-like PGBD | |
| 25 | >Braelyn gp42 | YP_010103934.1 | BE | BE1 | 471 | 50.5 | LysM-like PGBD | |
| 26 | >Egole gp45 | YP_010101465.1 | BE | BE1 | 474 | 50.7 | LysM-like PGBD | |
| 27 | >Tribute gp42 | QGH78234.1 | BE | BE1 | 474 | 50.5 | LysM-like PGBD | PGRP like amidase |
| 28 | >Sushi23 gp44 | ASR76476.1 | BE | BE1 | 473 | 50.7 | LysM-like PGBD | |
| 29 | >Samisti12 gp44 | YP_009611482.1 | BE | BE1 | 474 | 50.8 | LysM-like PGBD | |
| 30 | >MulchMansion gp41 | QNO12465.1 | BE | BE1 | 340 | 36.7 | Endolysin | 30% Amidase (4BOL), 22-28% LysM (4B9H and 4S3K) |
| 31 | >LilMartin gp41 | QNN98291.1 | BE | BE1 | 340 | 36.7 | endolysin | See MulchMansion |
| 32 | >Bmoc gp42 | YP_010107442.1 | BE | BE1 | 340 | 36.8 | endolysin | |
| 33 | >Peebs gp43 | YP_009611255.1 | BE | BE1 | 474 | 50.8 | LysM-like PGBD | 29% Amidase (4BOL), 27% LysM (4UZ2) |
| 34 | >LukeCage gp45 | YP_009839969.1 | BE | BE2 | 331 | 36.0 | LysM-like PGBD | |
| 35 | >StarPlatinum gp45 | YP_009839482.1 | BE | BE2 | 331 | 36.0 | LysM-like PGBD | |
| 36 | >Yaboi gp46 | YP_009841182.1 | BE | BE2 | 297 | 32.1 | LysM-like PGBD | |
| 37 | >Genie2 gp45 | QAY08707.1 | BE | BE2 | 297 | 32.1 | LysM-like PGBD | |
| 38 | >BoomerJR gp46 | QAY12697.1 | BE | BE2 | 297 | 32.1 | LysM-like PGBD | |
| 39 | >Wofford gp43 | YP_009839732.1 | BE | BE2 | 339 | 37.0 | LysM-like PGBD | |
| 40 | >MindFlayer gp44 | QPL13684.1 | BE | BE2 | 293 | 31.6 | Endolysin | |
| 41 | >SparkleGoddess gp22 | AXH68737.1 | BK | BK1 | 338 | 36.4 | LysM-like endolysin | |
| 42 | >Limpid gp21 | QGH79354.1 | BK | BK1 | 338 | 36.3 | LysM-like endolysin | |
| 43 | >Gilson gp25 | YP_009842488.1 | BK | BK1 | 338 | 36.5 | LysM-like endolysin | |
| 44 | >Comrade gp22 | YP_009840815.1 | BK | BK1 | 337 | 36.3 | LysM-like endolysin | |
| 45 | >Annadreamy gp21 | YP_009838987.1 | BK | BK1 | 338 | 36.3 | LysM-like endolysin | |
| 46 | >Blueeyedbeauty gp23 | YP_009839218.1 | BK | BK1 | 338 | 36.4 | LysM-like endolysin | |
| 47 | >Beuffert gp20 | QOI67422.1 | BK | BK1 | 350 | 37.3 | LysM-like endolysin | Align with 5JCE (26% identity) |
| 48 | >Wakanda gp17 | QIN94010.1 | BK | BK2 | 338 | 36.4 | lysin, LysM-like | |

*(table continues)*

| No | Endolysins | Genbank/pham Accession no. | Phage Cluster | Subcluster | AA length | Mass (kDa) | Predicted Function (by SEA-PHAGES) | Notes |
|---|---|---|---|---|---|---|---|---|
| **Amidase-LysM** | | | **20.0%** | | | | | |
| 49 | >Muntaha gp17 | QIN94574.1 | BK | BK2 | 336 | 36.3 | lysin, LysM-like LysM-like | Amidase (Autolysin: 4KNK_A residue 46-160 aligned with Attoomi residue 10-135 with 99.13% probability, 19% identity) HHpred |
| 50 | >Attoomi gp20 | YP_009796801.1 | Singleton | - | 348 | 37.4 | endolysin | |
| **PGRP/Amidase-PG-bd-like** | | | **39.6%** | | | | | |
| 1 | >VWB gp22 | AAR29740.1 | BA | None | 275 | 29.5 | N/A | |
| 2 | >Mojorita gp21 | APD18698.1 | BC | BC1 | 260 | 28.0 | endolysin | |
| 3 | >Picard gp21 | YP_009788224.1 | BC | BC1 | 260 | 28.0 | endolysin | |
| 4 | >SV1 gp21 | AFU62161.1 | BC | BC1 | 315 | 33.4 | putative endolysin | |
| 5 | >ToastyFinz gp21 | ARB11443.1 | BC | BC1 | 355 | 37.8 | endolysin | |
| 6 | >Raleigh gp22 | APD18773.1 | BC | BC2 | 360 | 37.8 | endolysin | |
| 7 | >Darolandstone gp22 | YP_009812933.1 | BC | BC2 | 361 | 37.5 | lysin A | |
| 8 | >Austintatious gp21 | YP_009819793.1 | BC | BC3 | 313 | 33.2 | endolysin | |
| 9 | >Ididsumtinwong gp21 | YP_009788115.1 | BC | BC3 | 313 | 33.2 | endolysin | |
| 10 | >PapayaSalad gp21 | YP_009788170.1 | BC | BC3 | 313 | 33.2 | endolysin | |
| 11 | >Bioscum gp21 | APD18718.1 | BC | BC3 | 313 | 33.3 | endolysin | |
| 12 | >Esperer gp26 | ATE85206.1 | BD | BD1 | 317 | 33.8 | endolysin | |
| 13 | >BryanRecycles gp27 | ATE84980.1 | BD | BD1 | 313 | 33.2 | endolysin | |
| 14 | >Oliynyk gp27 | ATE85357.1 | BD | BD1 | 313 | 33.2 | endolysin | |
| 15 | >BeardedLady gp27 | ATE84905.1 | BD | BD1 | 317 | 33.8 | endolysin | |
| 16 | >Jash gp27 | ATE85330.1 | BD | BD1 | 313 | 33.2 | endolysin | |
| 17 | >Nanodon gp28 | YP_009287812.1 | BD | BD1 | 314 | 32.9 | endolysin | |
| 18 | >Aaronocolus gp 26 | YP_009616451.1 | BD | BD1 | 317 | 33.8 | endolysin | |
| 19 | >Izzy gp27 | YP_009215405.1 | BD | BD1 | 313 | 33.2 | endolysin | |
| 20 | >Hydra gp28 | AKY03559.1 | BD | BD1 | 317 | 33.8 | LysM | |
| 21 | >Lannister gp28 | YP_009200968.1 | BD | BD1 | 317 | 33.6 | endolysin | |
| 22 | >Caliburn gp26 | YP_009207123.1 | BD | BD1 | 317 | 33.8 | endolysin | |
| 23 | >Phettuccine gp26 | QGJ91543.1 | BD | BD1 | 317 | 33.8 | lysin A | same as esperer |

*(table continues)*

| No | Endolysins | Genbank/pham Accession no. | Phage Cluster | Subcluster | AA length | Mass (kDa) | Predicted Function (by SEA-PHAGES) | Notes |
|---|---|---|---|---|---|---|---|---|
| **PGRP/Amidase-PG-bd-like** | | | **39.6%** | | | | | |
| 24 | >Rusticus gp27 | QDK03958.1 | BD | BD1 | 313 | 33.2 | LysM-like endolysin | same as esperer |
| 25 | >Leviticus gp26 | QDK03391.1 | BD | BD1 | 317 | 33.8 | LysM-like endolysin | PRPG 34%, AmpDh2 21% |
| 26 | >Nerdos gp26 | QAY17844.1 | BD | BD1 | 317 | 33.8 | LysM-like endolysin | same as esperer |
| 27 | >Indigo gp25 | QAY17301.1 | BD | BD1 | 317 | 33.8 | LysM-like endolysin | |
| 28 | >Bovely gp26 | QAY17229.1 | BD | BD1 | 317 | 33.8 | LysM-like endolysin | Amidase-lysM-PG-bdlike, PRPG 30%, AmpDh2 22% high coverage. |
| 29 | >Eddasa gp27 | AWN07470.1 | BD | BD1 | 313 | 33.2 | LysM-like endolysin | |
| 30 | >Ozzie gp26 | ATE85431.1 | BD | BD1 | 317 | 33.8 | endolysin | same as esperer |
| 31 | >Paedore gp26 | YP_010055892.1 | BD | BD2 | 358 | 37.6 | Lysin | |
| 32 | >R4 gp26 | AFU62079.1 | BD | BD2 | 356 | 37.1 | putative endolysin | same as ELB20 |
| 33 | >ELB20 gp25 | AFO10891.1 | BD | BD2 | 356 | 37.1 | N/A | |
| 34 | >Hank144 gp27 | YP_010055079.1 | BD | BD2 | 350 | 37.4 | Lysin A | |
| 35 | >Tefunt gp26 | YP_010055394.1 | BD | BD2 | 317 | 34.0 | Lysin | |
| 36 | >Diane gp26 | YP_010055236.1 | BD | BD2 | 317 | 34.0 | Lysin | |
| 37 | >Haizum gp26 | AXH70230.1 | BD | BD2 | 317 | 34.1 | Lysin A | |
| 38 | >Animus gp27 | QFG10695.1 | BD | BD2 | 358 | 38.1 | Lysin | |
| 39 | >Janus gp27 | QAY15931.1 | BD | BD2 | 358 | 38.1 | Lysin A | |
| 40 | >Nishikigoi gp26 | QAY15767.1 | BD | BD2 | 317 | 34.1 | Lysin A | |
| 41 | >Amethyst gp26 | YP_010055315.1 | BD | BD2 | 316 | 34.0 | Lysin | |
| 42 | >SqueakyClean gp27 | ATI18890.1 | BD | BD2 | 358 | 38.1 | Lysin | |
| 43 | >phiCAM gp28 | YP_009592107.1 | BD | BD3 | 226 | 24.3 | N/A | |
| 44 | >Yosif gp28 | YP_010054670.1 | BD | BD3 | 317 | 34.1 | Lysin | |
| 45 | >Verse gp27 | AKY03857.1 | BD | BD3 | 323 | 34.6 | endolysin | See Amela |
| 46 | >Amela gp27 | AKY03782.1 | BD | BD3 | 323 | 34.5 | endolysin | 34% PRPG (4C8I)/Amidase (2F2L), 20%AmpDh2 (4BOL) |
| 47 | >phiHau3 gp28 | YP_006906203.1 | BD | BD4 | 350 | 37.2 | putative endolysin | |
| 48 | >Urza gp28 | QFG10491.1 | BD | BD6 | 328 | 34.9 | lysin A | |
| 49 | >Celia gp28 | YP_010054592.1 | BD | BD6 | 328 | 34.9 | LysM-like endolysin | |
| 50 | >Daubenski gp45 | YP_010104808.1 | BE | BE1 | 298 | 31.9 | LysM-like PGBD | |
| 51 | >Wipeout gp44 | QGH74290.1 | BE | BE2 | 293 | 31.6 | LysM-like PGBD | |
| 52 | >TomSawyer gp44 | QGH78931.1 | BE | BE2 | 293 | 31.6 | LysM-like PGBD | |

*(table continues)*

| No | Endolysins | Genbank/pham Accession no. | Phage Cluster | Subcluster | AA length | Mass (kDa) | Predicted Function (by SEA-PHAGES) | Notes |
|---|---|---|---|---|---|---|---|---|
| PGRP/Amidase-PG-bd-like | | | 39.6% | | | | | |
| 53 | >Starbow gp44 | AXH66553.1 | BE | BE2 | 293 | 31.6 | LysM-like PGBD | |
| 54 | >Birchlyn gp43 | QDF17219.1 | BE | BE2 | 293 | 31.6 | LysM-like PGBD | |
| 55 | >Karimac gp45 | YP_009840217.1 | BE | BE2 | 293 | 31.6 | LysM-like PGBD | |
| 56 | >IchabodCrane gp43 | QFP97359.1 | BE | BE2 | 293 | 31.6 | LysM-like PGBD | |
| 57 | >Bordeaux gp44 | QGH79816.1 | BE | BE2 | 293 | 31.6 | LysM-like PGBD | Same as Karimac gp45 |
| 58 | >HaugeAnator gp2 | AUG87329.1 | BF | None | 253 | 28.0 | Lysin A | |
| 59 | >ZooBear gp2 | AUG87585.1 | BF | None | 253 | 28.0 | Lysin A | |
| 60 | >ToriToki gp 2 | AUG87521.1 | BF | None | 254 | 28.1 | Lysin A | |
| 61 | >Romero gp2 | AUG87457.1 | BF | None | 254 | 28.1 | Lysin A | |
| 62 | >Percastrophe gp2 | AUG87393.1 | BF | None | 254 | 28.1 | Lysin A | |
| 63 | >Olicious gp2 | AZF95812.1 | BF | None | 253 | 28.0 | Lysin A | |
| 64 | >Immanuel3 gp2 | YP_009836121.1 | BF | None | 254 | 28.1 | Lysin A | |
| 65 | >Geostin gp2 | QEA11225.1 | BF | None | 257 | 27.9 | Lysin A | Same as Fabian |
| 66 | >Fabian gp2 | QFP94721.1 | BF | None | 255 | 27.7 | Lysin A | |
| 67 | >FlowerPower gp2 | YP_009838059.1 | BF | None | 257 | 27.9 | Lysin A | Same as Fabian |
| 68 | >WRightOn gp3 | YP_009835995.1 | BF | None | 259 | 28.3 | Lysin A | |
| 69 | >Manuel gp2 | YP_009836058.1 | BF | None | 233 | 25.5 | Lysin A | |
| 70 | >Salete gp27 | AWN08458.1 | BG | None | 420 | 44.1 | Lysin | |
| 71 | >BayC gp27 | AWN08387.1 | BG | None | 420 | 44.1 | Lysin | |
| 72 | >Abt2graduatex2 gp28 | ATN93711.1 | BG | None | 425 | 44.8 | Lysin | |
| 73 | >BabyGotBac gp27 | APZ82195.1 | BG | None | 420 | 44.1 | Lysin | |
| 74 | >Maih gp27 | ALY07277.1 | BG | None | 420 | 44.1 | Lysin | |
| 75 | >Xkcd426 gp33 | AMD42774.1 | BG | None | 335 | 35.6 | Lysin | |
| 76 | >TP1604 gp27 | AKA61765.1 | BG | None | 420 | 44.1 | Lysin | |
| 77 | >YDN12 gp29 | AKA61696.1 | BG | None | 422 | 44.4 | Lysin | |
| 78 | >Mischief19 gp36 | QBZ73521.1 | BG | None | 321 | 34.2 | Lysin A | |
| 79 | >Dubu gp18 | QDH92123.1 | BJ | None | 312 | 34.1 | lysin A, N-acetylmuramoyl-L-alanine amidase domain | |
| 80 | >phiSASD1 gp40 | YP_003714747.1 | BJ | None | 278 | 29.6 | endolysin | 22% Amidase (1YB0), 27% PG-bd-like 5NM7 |
| 81 | >Moab gp22 | QIQ62907.1 | BK | BK1 | 299 | 31.9 | Lysin A | same as IchabodCrane gp43 |
| 82 | >Satis gp118 | AXH66279.1 | BM | None | 361 | 38.0 | lysin A | AmphD2, 30% identity with high coverage |
| 83 | >JustBecause gp116 | AYD81285.1 | BM | None | 361 | 38.2 | lysin A | same as Kradal |

*(table continues)*

| No | Endolysins | Genbank/pham Accession no. | Phage Cluster | Subcluster | AA length | Mass (kDa) | Predicted Function (by SEA-PHAGES) | Notes |
|---|---|---|---|---|---|---|---|---|
| **PGRP/Amidase-PG-bd-like** | | | **39.6%** | | | | | |
| 84 | >Kradal gp118 | QBZ72016.1 | BM | None | 361 | 38.0 | lysin A | looks like amidase-lysm, 28% identity to AmpDh2 with high coverage |
| 85 | >Yara gp25 | AVP41359.1 | BN | None | 307 | 33.9 | lysin | |
| 86 | >Gibson gp28 | QAX92974.1 | BN | None | 309 | 32.8 | endolysin | |
| 87 | >Wentworth gp29 | AVP41468.1 | BN | None | 305 | 32.6 | lysin | |
| 88 | >Forthebois gp35 | YP_010084058.1 | BO | None | 258 | 27.0 | LysM-like endolysin | |
| 89 | >WheeHeim gp36 | YP_010084094.1 | BO | None | 261 | 27.3 | LysM-like endolysin | |
| 90 | >Hiyaa gp30 | YP_009818466.1 | BQ | None | 324 | 35.9 | LysM-like endolysin | |
| 91 | >BRock gp39 | YP_009831765.1 | Singleton | None | 310 | 34.5 | N/A | Amidase (AmiE 3LAT 20%), PG-bd-like 31% 5TV7 |
| 92 | >Chymera gp20 | AMS01579.1 | Singleton | None | 291 | 31.6 | Lysin | |
| 93 | >Zuko gp35 | QEQ93613.1 | Singleton | None | 317 | 34.4 | lysin A, N-acetylmuramoyl-L-alanine amidase domain | |
| 94 | >pZL12 gp44 | ACX71121.1 | Singleton | None | 312 | 33.0 | N/A | |
| 95 | >phiSAV gp25 | BAC73210.1 | Singleton | None | 257 | 27.7 | N/A | |
| 96 | >Gilgamesh gp88 | QFG13280.1 | Singleton | None | 351 | 37.4 | lysin A, N-acetylmuramoyl-L-alanine amidase domain | |
| 97 | >Kromp gp25 | AYD81626.1 | Singleton | None | 308 | 33.1 | LysM-like endolysin | |
| 98 | >Ibantik gp85 | AWN05307.1 | Singleton | None | 351 | 37.4 | Lysin A | |
| 99 | >mu1/6 gp19 | YP_579222.1 | Singleton | None | 393 | 42.1 | N/A | Amidase aligned with AmpDh2 and 3 (20% identity, high coverage) Pgbdlike from peptidase (1LBU, 30% identity) |
| **Amidase-X** | | | **1.2%** | | | | | |
| 1 | >Kardashian gp4 | QEQ93869.1 | BI | BI6 | 286 | 31.7 | Lysin A | 26% XylA (3HMA), 38% incomplete CW-7 (5I8L) but CW-7 doesn't align very well. |
| 2 | >SendItCS gp4 | AWN06107.1 | BI | BI4 | 280 | 30.9 | N/A | 23% amidase (AmpDh3 4BXD), 40% incomplete CW-7 (5I8L) |
| 3 | >Rainydai gp4 | AWN05870.1 | BI | BI4 | 280 | 30.9 | N/A | see SendItCS gp4 |
| **Amidase-Transglycosylase** | | | **2.0%** | | | | | |
| 1 | >Warpy gp48 | ASN73122.1 | BE | BE1 | 472 | 50.4 | LysM-like PGBD | See evy gp45 |
| 2 | >Evy gp45 | YP_010103421.1 | BE | BE1 | 473 | 50.5 | LysM-like PGBD | 27% amidase (4BOL), 32% transglycosylase active site (1GBS) |
| 3 | >Jay2Jay gp49 | AIW02547.1 | BE | BE1 | 474 | 50.8 | LysM-like PGBD | See evy gp45 |
| 4 | >Circinus gp19 | QBZ72303.1 | BK | BK2 | 416 | 44.6 | LysM-like endolysin | |
| 5 | >BillNye gp17 | YP_009622595.1 | BK | BK2 | 418 | 44.7 | LysM-like endolysin | |

*(table continues)*

| No | Endolysins | Genbank/pham Accession no. | Phage Cluster | Subcluster | AA length | Mass (kDa) | Predicted Function (by SEA-PHAGES) | Notes |
|---|---|---|---|---|---|---|---|---|
| **CHAP-LysM** | | | **0.4%** | | | | | |
| 1 | >Sros11 gp23 | pham 97340 | BA | None | 310 | 33.0 | N/A | |
| **CHAP-PG-bd-like** | | | **13.2%** | | | | | |
| 1 | >phiC31 gp50 | CAA07120.2 | BB | BB1 | 362 | 38.4 | N/A | 26% to C-ter of 4HPE, PG-bd-like 30%to 5TV7 |
| 2 | >phiBT1 gp50 | CAD80117.1 | BB | BB1 | 278 | 29.8 | N/A | |
| 3 | >Vash gp19 | QAX93275.1 | BB | BB1 | 275 | 29.2 | lysin A | |
| 4 | >Lilbooboo gp19 | YP_009819898.1 | BB | BB1 | 273 | 29.0 | lysin A | See Vash |
| 5 | >Euratis gp19 | QAX94014.1 | BB | BB1 | 275 | 29.5 | lysin A | 20-23% CHAP (4HPE and 5UDM), 28% PG-bd-like (5TV7) |
| 6 | >Shawty gp19 | QAY26944.1 | BB | BB1 | 280 | 29.8 | lysin A | |
| 7 | >TG1 gp19 | AFU62214.1 | BB | BB1 | 276 | 29.8 | putative endolysin | |
| 8 | >RemusLoopin gp19 | QBZ73333.1 | BB | BB2 | 270 | 29.4 | lysin A | |
| 9 | >Heather gp19 | QBZ73389.1 | BB | BB2 | 279 | 31.3 | lysin A | |
| 10 | >Sebastisaurus gp19 | QAX95007.1 | BB | BB2 | 275 | 29.5 | lysin A | |
| 11 | >Thestral gp27 | YP_010055810.1 | BD | BD2 | 363 | 39.0 | Lysin A | |
| 12 | >Omar gp27 | AUG87211.1 | BD | BD2 | 286 | 31.2 | Lysin | |
| 13 | >TinaBelcher gp26 | QAY15849. | BD | BD2 | 363 | 39.1 | Lysin A | |
| 14 | >Bowden gp27 | QAY15685.1 | BD | BD2 | 369 | 39.6 | Lysin A | |
| 15 | >Alvy gp28 | YP_010055722.1 | BD | BD2 | 370 | 40.0 | Lysin A | |
| 16 | >Daudau gp26 | YP_010055638.1 | BD | BD2 | 280 | 29.8 | Lysin | |
| 17 | >TrvxScott gp25 | AXQ62351.1 | BD | BD2 | 368 | 39.3 | Lysin A | |
| 18 | >BartholomewSD gp28 | QAX95478.1 | BD | BD2 | 370 | 40.0 | Lysin A | |
| 19 | >Caelum gp27 | YP_010055476.1 | BD | BD2 | 285 | 30.8 | Lysin A | |
| 20 | >Alsaber gp28 | YP_010056359.1 | BD | BD3 | 285 | 31.0 | N/A | |
| 21 | >Saftant gp27 | YP_010056207.1 | BD | BD3 | 288 | 31.0 | Lysin A | |
| 22 | >StrepC gp26 | pham 97340 | BD | BD5 | 276 | 30.1 | N/A | |
| 23 | >Araceli gp35 | QFG07849.1 | BH | None | 248 | 26.8 | endolysin | |
| 24 | >Intolerant gp34 | QFG07929.1 | BH | None | 252 | 27.1 | endolysin | |
| 25 | >Microdon gp34 | AYD86781.1 | BH | None | 252 | 27.1 | LysM-like endolysin | |
| 26 | >LazerLemon gp35 | AWY07518.1 | BH | None | 252 | 27.5 | endolysin | |
| 27 | >Crosby gp35 | AXH49423.1 | BH | None | 252 | 27.2 | lysin A | |
| 28 | >UNTPL gp35 | AWY07436.1 | BH | None | 252 | 27.0 | endolysin | |

*(table continues)*

| No | Endolysins | Genbank/pham Accession no. | Phage Cluster | Subcluster | AA length | Mass (kDa) | Predicted Function (by SEA-PHAGES) | Notes |
|---|---|---|---|---|---|---|---|---|
| CHAP-PG-bd-like | | | 13.2% | | | | | |
| 29 | >Henoccus gp35 | AWY07273.1 | BH | None | 250 | 26.9 | endolysin | |
| 30 | >JackieB gp34 | AWY07353.1 | BH | None | 248 | 26.8 | endolysin | |
| 31 | >Nesbitt gp21 | AVO22278.1 | BL | None | 284 | 30.6 | lysin | |
| 32 | >Rowa gp21 | YP_009796855.1 | BL | None | 284 | 30.1 | lysin | |
| 33 | >AbbeyMikolon gp21 | YP_009796743.1 | BL | None | 277 | 29.8 | lysin | |
| Zinc peptidase-CW-7 | | | 10.8% | | | | | |
| 1 | >FrodoSwaggins gp4 | QGJ96544.1 | BI | BI1 | 291 | 32.4 | endolysin | |
| 2 | >FidgetOrca gp5 | QGJ96704.1 | BI | BI1 | 289 | 32.2 | endolysin | |
| 3 | >Meibysrarus gp5 | QEQ93953.1 | BI | BI1 | 289 | 32.2 | endolysin | |
| 4 | >Jaylociraptor gp5 | QEQ93698.1 | BI | BI1 | 289 | 32.2 | endolysin | |
| 5 | >Hoshi gp5 | QEQ94222.1 | BI | BI1 | 289 | 32.2 | endolysin | |
| 6 | >GirlPower gp4 | QEQ93503.1 | BI | BI1 | 288 | 32.0 | endolysin | |
| 7 | >CherryBlossom gp5 | QEQ93784.1 | BI | BI1 | 289 | 32.0 | endolysin | |
| 8 | >Esketit gp5 | QDM56506.1 | BI | BI1 | 289 | 32.2 | lysin A | |
| 9 | >Soshi gp4 | QEQ94617.1 | BI | BI1 | 291 | 32.4 | endolysin | |
| 10 | >Popy gp4 | QAY17038.1 | BI | BI1 | 291 | 32.4 | endolysin | |
| 11 | >Namo gp5 | QAY16303.1 | BI | BI1 | 289 | 32.2 | endolysin | |
| 12 | >Madamato gp5 | QAY17123.1 | BI | BI1 | 289 | 32.1 | endolysin | |
| 13 | >Spectropatronm gp5 | ASU04001.1 | BI | BI1 | 289 | 32.2 | endolysin | |
| 14 | >DrGrey gp4 | YP_009612534.1 | BI | BI1 | 291 | 32.4 | endolysin | |
| 15 | >IceWarrior gp5 | QAY16217.1 | BI | BI1 | 289 | 32.2 | endolysin | |
| 16 | >Rima gp5 | AOZ64959.1 | BI | BI1 | 289 | 32.2 | endolysin | |
| 17 | >OlympicHelado gp5 | AOZ64870.1 | BI | BI1 | 289 | 32.2 | endolysin | |
| 18 | >Maya gp5 | QNN98170.1 | BI | BI1 | 289 | 32.2 | endolysin | |
| 19 | >TieDye gp4 | QNN99480.1 | BI | BI1 | 289 | 32.2 | endolysin | |
| 20 | >PherryCruz gp3 | QBZ73430.1 | BI | BI2 | 294 | 32.9 | N/A | |
| 21 | >RavenPuff gp3 | AWN05960.1 | BI | BI2 | 294 | 32.9 | N/A | |
| 22 | >Moozy gp4 | AWN05439.1 | BI | BI2 | 294 | 32.8 | N/A | |
| 23 | >HotFries gp3 | AWN05171.1 | BI | BI2 | 294 | 32.8 | N/A | |

*(table continues)*

| No | Endolysins | Genbank/pham Accession no. | Phage Cluster | Subcluster | AA length | Mass (kDa) | Predicted Function (by SEA-PHAGES) | Notes |
|---|---|---|---|---|---|---|---|---|
| **Zinc peptidase-CW-7** | | | **10.8%** | | | | | |
| 24 | >Scap1 gp4 | ATN93653.1 | BI | BI2 | 289 | 32.2 | endolysin | |
| 25 | >LibertyBell gp4 | AXQ61245.1 | BI | BI3 | 279 | 31.4 | endolysin | |
| 26 | >Bing gp4 | YP_009622802.1 | BI | BI5 | 290 | 32.1 | endolysin | |
| 27 | >RosaAsantewaa gp4 | QBZ73568.1 | Singleton | None | 287 | 32.0 | N/A | |
| **Glycosyl hydrolase** | | | **0.4%** | | | | | |
| 1 | >Shyg gp25 | pham 4657 | BC | BC4 | 304 | 32.6 | N/A | |
| **X-Transglycosylase (SAR)** | | | **12.4%** | | | | | |
| 1 | >Evy gp40 | YP_010103416.1 | BE | BE1 | 174 | 19.0 | Endolysin | |
| 2 | >Daubenski gp41 | YP_010104804.1 | BE | BE1 | 178 | 19.2 | Endolysin | |
| 3 | >Braelyn gp37 | YP_010103929.1 | BE | BE1 | 179 | 19.8 | Endolysin | See NootNoot |
| 4 | >Teutsch gp39 | QAX95775.1 | BE | BE1 | 182 | 19.7 | Endolysin | |
| 5 | >Egole gp40 | YP_010101461.1 | BE | BE1 | 184 | 19.8 | Endolysin | |
| 6 | >Tribute gp38 | QGH78230.1 | BE | BE1 | 182 | 19.7 | Endolysin | |
| 7 | >Mildred21 gp38 | YP_009610579.1 | BE | BE1 | 197 | 21.6 | Endolysin | 24% transglycosylase (1GSA, 1SLY), no N-term predicted |
| 8 | >NootNoot gp34 | YP_009610809.1 | BE | BE1 | 179 | 19.7 | Endolysin | |
| 9 | >Paradiddles gp34 | YP_009611030.1 | BE | BE1 | 179 | 19.8 | Endolysin | see NootNoot |
| 10 | >Warpy gp42 | ASN73116.1 | BE | BE1 | 178 | 19.4 | Endolysin | See Tribute |
| 11 | >Sushi23 gp40 | ASR76472.1 | BE | BE1 | 183 | 19.6 | Endolysin | see Teutsch gp39 |
| 12 | >Samisti12 gp39 | YP_009611477.1 | BE | BE1 | 182 | 19.7 | Endolysin | See Peebs |
| 13 | >Peebs gp39 | YP_009611251.1 | BE | BE1 | 182 | 19.7 | Endolysin | See Egole |
| 14 | >Jay2Jay gp42 | AIW02540.1 | BE | BE1 | 178 | 19.4 | Endolysin | See Warpy |
| 15 | >MulchMansion gp37 | QNO12461.1 | BE | BE1 | 178 | 19.2 | Hydrolase | |
| 16 | >LilMartin gp37 | QNN98287.1 | BE | BE1 | 178 | 19.2 | N/A | |
| 17 | >Bmoc gp37 | YP_010107437.1 | BE | BE1 | 175 | 18.8 | endolysin | 21% transglycosylase (4HPE,4OW1, 6CFC), no N-term predicted |
| 18 | >IchabodCrane gp38 | QFP97354.1 | BE | BE2 | 181 | 20.1 | Endolysin | See Karimac |
| 19 | >Bordeaux gp39 | QGH79811.1 | BE | BE2 | 181 | 20.0 | Endolysin | See Karimac |
| 20 | >Genie2 gp41 | QAY08702.1 | BE | BE2 | 181 | 20.2 | Endolysin | |
| 21 | >BoomerJR gp41 | QAY12692.1 | BE | BE2 | 181 | 20.2 | Endolysin | See Yaboi |
| 22 | >Yaboi gp41 | YP_009841177.1 | BE | BE2 | 181 | 20.2 | Endolysin | See Karimac |
| 23 | >Wipeout gp38 | QGH74284.1 | BE | BE2 | 181 | 20.0 | Endolysin | See Karimac |
| 24 | >TomSawyer gp39 | QGH78926.1 | BE | BE2 | 181 | 20.0 | Endolysin | See Karimac |
| 25 | >Wofford gp39 | YP_009839727.1 | BE | BE2 | 184 | 20.3 | Endolysin | 29% transglycosylase (1GSA), no N-term predicted. |

*(table continues)*

| No | Endolysins | Genbank/pham Accession no. | Phage Cluster | Subcluster | AA length | Mass (kDa) | Predicted Function (by SEA-PHAGES) | Notes |
|---|---|---|---|---|---|---|---|---|
| **X-Transglycosylase (SAR)** | | | **12.4%** | | | | | |
| 26 | >Starbow gp39 | AXH66548.1 | BE | BE2 | 181 | 20.1 | Endolysin | See Karimac |
| 27 | >LukeCage gp40 | YP_009839964.1 | BE | BE2 | 181 | 20.2 | Endolysin | See Karimac |
| 28 | >StarPlatinum gp40 | YP_009839477.1 | BE | BE2 | 181 | 20.1 | Endolysin | See Karimac |
| 29 | >Birchlyn gp38 | QDF17214.1 | BE | BE2 | 181 | 20.0 | Endolysin | See Karimac |
| 30 | >Karimac gp40 | YP_009840212.1 | BE | BE2 | 181 | 20.1 | Endolysin | |
| 31 | >MindFlayer gp39 | QPL13679.1 | BE | BE2 | 181 | 20.1 | Hydrolase | See Karimac |
